# Area and Geometry Dependence of Cell Migration in Asymmetric Two-State Micropatterns

**DOI:** 10.1101/815472

**Authors:** Alexandra Fink, David B. Brückner, Christoph Schreiber, Peter J. F. Röttgermann, Chase P. Broedersz, Joachim O. Rädler

## Abstract

Micro-structured surfaces provide a unique framework to probe cell migration and cytoskeletal dynamics in a standardized manner. Here, we report on the steady-state occupancy probability of cells in asymmetric two-state microstructures that consist of two fibronectin-coated adhesion sites connected by a thin guidance cue. In these dumbbell-like structures, cells transition between the two sites in a repeated and stochastic manner and average dwell times in the respective microenvironments are determined from the cell trajectories. We study the dynamics of human breast carcinoma cells (MDA-MB-231) in these microstructures as a function of area, shape and orientation of the adhesion sites. On square adhesive sites with different areas, we find that the occupancy probability ratio is directly proportional to the ratio of corresponding adhesion site areas. Sites of equal area but different shape lead to equal occupancy, if shapes are isotropic, e.g. squared or circular. In contrast, an asymmetry in the occupancy is induced by anisotropic shapes like rhombi, triangles or rectangles that enable motion in the direction perpendicular to the transition axis. Analysis of the 2D motion of cells between two rectangles with orthogonal orientation suggests that cellular transition rates depend on the cell polarisation induced by anisotropic micropatterns. Taken together, our results illustrate how two-state-micropatterns provide a dynamic migration assay with distinct dwell times and relative cell occupancy as readouts, which may potentially be useful to probe cell-microenvironment interactions.

## Introduction

In many fundamental biological processes like early development, cancer metastasis and inflammation, cells are guided by external cues. Understanding the factors and mechanisms that steer cells in tissue and in the extracellular matrix is still an important open biophysical challenge. Many cell-based assays provide platforms to quantify cell behaviour in purposefully designed settings. Well studied are the mechanisms of cell guidance by external cues, for example chemotaxis enabling cells to follow soluble chemical gradients, or durotaxis, in which a motile cell is guided by mechanical stiffness gradients in the surrounding matrix (1–5). Likewise, gradients of surface bound integrin ligands and differences in extracellular matrix protein density can guide cell migration in two dimensions (6–8). Cellular preferences or decision making is typically probed on the population level. In such experiments, cell-substrate interactions are tuned over a large surface area and the distribution of cells is measured as endpoint readout rather than in a time-resolved manner (9–15). A more direct way to probe cell decision making in the presence of guidance cues is to provide assays where single cells are given the choice to spread on one of two different kinds of surfaces. In neuronal research, biochemical assays comprising adjacent lines of different proteins are known to study the preferential alignment and adhesion of neurons (16, 17). Also, the response of sensory neurites and growth cones to different substrates and patterns of different proteins has been studied (18, 19). Thus, micropatterning has advanced as a tool for single cell adhesion and migration studies providing standardized microenvironments.

Various micro-structuring techniques enable the spatially controlled deposition of proteins in defined areas (20–23). Cells seeded on micropatterns spread and adapt to the shape and size of the adhesive area (24). Regular lattices of micropatterns have been used to capture and strain single cells for a multitude of studies, including spindle-orientation (25, 26), the influence of cell shape on cell polarization (27–31), and to monitor cell responses over time (32, 33). For the investigation of cell migration dynamics, micropatterned lanes have proven useful as they confine cell motion to one dimension (34–37). Variations of microlanes with symmetry-breaking triangular or teardrop-shaped patterns demonstrate control over the direction of cell migration by geometry (37–41). All these studies benefit from highly reproducible microenvironments and extensive single cell statistics.

However, few experiments exist that combine geometric adhesion sites with dynamic migration experiments. Often the dynamics of the decision making is not monitored and it remains unknown whether in repeated trials a single cell would respond in the same manner. This is all the more important as cellular heterogeneity is understood to be an intrinsic property of cell cultures (42). It is hence our goal to design confining geometries such that single migrating cells repeatedly explore their micro-environments. The general question we like to address is how cellular dynamics is biased by the geometry of their respective micro-environment.

Recently, we reported on a novel type of confined migration of cells in two-state micropatterns (43). The symmetric micropatterns consist of two approximately cell-sized adhesive sites, connected by a thin bridge. On these dumbbell-shaped structures, cells transition between the two sites in a repeated and stochastic manner. The theoretical analysis decomposed cell motion in this particular confinement into distinct deterministic and stochastic components and demonstrated that different cell lines can exhibit distinct nonlinear deterministic dynamics. This setup provides a unique platform to study confined cell migration in a controlled and standardized assay, and thus forms an ideal system to study how cells respond to adhesion site geometries.

In this study, we explore the dwell times and steady-state distribution of cells on two-state micropatterns with adhesive sites that feature different geometric properties. This system represents a simple implementation of a confining microenvironment, which is capable to probe the preference of cells for defined surface geometries. The micropatterns consist of two fibronectin-coated adhesion sites that are connected by a thin fibronectin-coated stripe. In these structures, cells transition repeatedly between the two adhesion sites. We monitor the trajectories of the fluorescently labelled nuclei and quantify the occupation probabilities on each adhesion site. We show that the relative dwell times between subsequent transitions on differently sized square adhesion sites scales approximately linearly with the ratio of the adhesion areas. Pairs of sites with equal area but different isotropic geometries, such as squares and circles, result in equal distributions indicating a weak effect of site shapes on dwell times. However, patterns that include anisotropic sites, e.g. squares versus rhombi or triangles show skewed occupancies. We attribute this effect to induced cell polarization and demonstrate this in the case of equally sized rectangular adhesion sites arranged in different orientations with respect to the connecting bridge. In this case, we observe a preferred occupancy in the rectangle that is oriented perpendicular to the direction of transitions. Our observations are consistent with a simple kinetic rate model and we discuss possible applications of dynamic assays to measure relative affinities of cells in artificially designed asymmetric microenvironments.

## Materials and Methods

### Micropatterning and Sample Preparation

We use microscale plasma-induced protein patterning (44). Briefly, silicon masters with the desired shapes are prepared using photolithography. Polydimethylsiloxane (PDMS) monomer and crosslinker (DC 184 elastomer kit, Dow Corning, Midland, Michigan) are mixed in a 10:1 ratio and subsequently poured onto the silicon wafer. After degassing the PDMS, it is cured overnight at 50 °C.

PDMS stamps are placed, with the features facing down, in an ibidi µ-dish (ibidi GmbH, Martinsried, Germany): The dish with the stamps is then exposed to oxygen plasma. For background passivation, a drop of 2 mg ml^−1^ PLL(20)-g[3.5]-PEG(2) (SuSoS AG, Dübendorf, Switzerland) solution is added and left to incubate for 25 min. After rinsing the sample, the stamps are removed, and the sample is incubated with a 50 µg ml^−1^ human fibronectin (YO Proteins, Ronninge, Sweden) solution for 50 min. After final rinsing, samples are stored in PBS at 4°C.

### Micropattern Design

The two-state micropatterns have square adhesion sites of edge lengths between (27.3 ± 0.4) µm to (42.2 ± 0.5) µm. Adhesive area dimensions were chosen such as to provide enough adhesive area for cells to be able to fully occupy them. A limit was found for adhesion sites smaller than approximately 27 µm, where cells would still transition between adhesion sites but partially remain on the bridge and the larger adhesion site. No adhesion sites with edge lengths larger than 42.2 µm were used.

Square-circle micropatterns were designed to have the circular sites either probe a same-area or a same-perimeter case with respect to the square adhesion sites. Micropatterns with triangles are designed to offer adhesion sites of equal areas and also to both have a right angle pointing towards the bridge. Rectangular adhesion sites have equal areas and were designed to offer similar adhesion site areas to the symmetric square-square setup. The aspect ratio of rectangles was chosen to be 2:1. For all geometries, a similar bridge length was used. The influence of bridge length was also tested for rectangular patterns. Details of that and exact measures for all patterns, as well as cell statistics, can be found in the Supplementary Sections 1 and 5.

Errors given for the pattern dimensions are weighted standard deviations to account for the variation in statistics gained in different experiments. The dimensions of final protein patterns are subject to experiment-to-experiment and wafer-to-wafer variations. Experiment-to-experiment variability is mainly due to the intrinsic variance of the manual stamping process. Also, measurement uncertainty is added due to the limited resolution of images.

### Cell Culture

MDA-MB-231 human breast carcinoma epithelial cells (DSMZ, Braunschweig, Germany) are cultured in Minimum Essential Medium (MEM, c.c. pro, Oberdorla, Germany) with 10% FBS (Gibco, Paisley, United Kingdom) and 2mM L-Glutamine (c.c. pro). Cells are grown in an atmosphere with 5% CO_2_ and at 37°C up to 70-90% confluence. For splitting, cells are rinsed with PBS and subsequently trypsinised for 3 min. For experiments, the trypsinised cell solution is centrifuged at 1000 rcf for 3 min, the cell pellet is re-suspended in MEM and approximately 10,000 cells are seeded per µ-dish. Cells are left to adhere for 4h in the incubator, before the medium is exchanged to L-15 medium (containing L-Glutamine (Gibco, Grand Island, New York), and supplemented with 10% FCS). The L-15 also contains 25 nM Hoechst 33342 (Invitrogen, Eugene, Oregon) for the staining of cell nuclei.

### Microscopy and Cell Tracking

Measurements of up to 50h are performed in time-lapse mode, either on an IMIC digital microscope (TILL Photonics, Kaufbeuren, Germany) or on a Nikon Eclipse Ti microscope using a 10x objective. To keep samples at 37°C throughout the measurement, dishes are placed in a heated chamber (ibidi GmbH or Okolab, Pazzuoli, Italy). Brightfield and DAPI-images are acquired every 10 min. For analysis, a band pass filter is applied to the fluorescence images and a threshold is used to binarise the images using ImageJ (45). This allows subsequent automated tracking of the centre-of-mass of the cell nuclei by ImageJ’s Analyze Particles plugin. The two-state patterns are visible in the brightfield images and are used to manually determine the reference boundary of a pattern, defined as edge of the left adhesive island, for each single cell. Using this value it is possible to convert nuclear coordinates to absolute positions. All further analysis is performed in MATLAB (The MathWorks, Natick, Massachusetts).

For every geometry, at least 46 cells, chosen from a minimum of three experiments, were analysed. The criteria for the determination of suitable cells, as well as the number of analysed cells per geometry, are detailed in Supplementary Section S1.

### LifeAct-GFP transfection

Approximately 12,500 MDA-MB-231 cells are seeded in patterned µ-dishes and left to adhere overnight. Cells are cultured in MEM including Glutamax (Gibco, Paisley, United Kingdom) supplemented with 10% FCS.

For life-cell actin imaging, 500ng LifeAct-GFP mRNA (in-house prepared) is resuspended in OptiMEM (Gibco, Grand Island, New York) to a final volume of 150µl. In parallel, a mix of 1.25 µl Lipofectamine 2000 (Invitrogen, Carlsbad, California) and 123.75 µl OptiMEM is prepared. Then, the mRNA solution is added to the Lipofectamine-mix, and left to incubate for 20 minutes at room temperature. Before adding the transfection mix, cells are rinsed once with PBS. Cells are incubated with the transfection mix for at least 3h, before the mix is replaced by L-15 medium. Cells are imaged every 10 minutes on the Nikon Ti Eclipse microscope using a 60x oil-immersion objective.

### Dwell Times and Occupation Probabilities

For dwell time calculation, the trajectories of the nuclei are binarised into two states, left and right of the middle of the connecting bridge. Furthermore, in order to avoid effects of a finite observation time window, all analysis is performed on cropped trajectories, starting from the time point of the first transition and ending after the last fully observed dwell time. The middle of the bridge is determined by adding the sum of the mean left adhesion site edge length and half of the mean bridge length for each experiment to the individually determined left border reference points. The time spent in either state in between transitions over the middle of the bridge, is the dwell time τ.

To assess the relative occupation probabilities, we calculate the sum of dwell times, *τ*_*i*_, on adhesion site i divided by the total observation time, T_tot_ of all trajectories, which can also be expressed by the number of stays N_i_ and the mean stay time, ⟨*τ*_*i*_⟩, divided by total time:

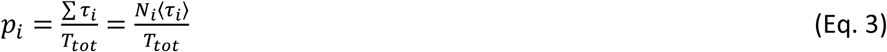

The ratio of the observed left/right mean dwell times ⟨*τ*_*left*_⟩/ ⟨*τ*_*right*_⟩ thus converges to the ratio of probabilities p_left_/p_right_, if the number of total stays is large, and thus N_left_≈N_right_.

To correct for unequal adhesion site areas in the cases where adhesion sites were designed to have the same area but were found to deviate from the expected areas, we normalise the occupation probabilities by adhesion site areas A:

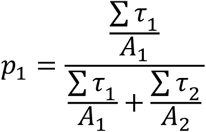

### Survival Probability Functions

The survival probability distribution S(t) is defined as

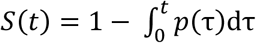

where p(τ) is the probability distribution of dwell times for all cells evaluated on a certain adhesion site.

### Error Analysis

We employ a bootstrapping procedure to estimate errors (43, 46). We generate a large number of realisations of a given dataset *D* = {*X*_1_,…,*X*_*N*_} with entries by randomly sampling the dataset’s entries with replacement. The mean ⟨*X*⟩_realisation_ is calculated for each realisation. The standard deviation of all ⟨*X*⟩_realisation_ gives the estimated error in the mean.

The same procedure is carried out for small groups of subsequent entries of the dataset to account for correlations. For each window size, 50000 realisations are generated. We limit the window size to a maximum value of 60. The maximum value of the standard deviation calculated for each window size is our final bootstrap error.

## Results

### Cells repeatedly transition between adhesion sites on asymmetric two-state micropatterns

To study how the shape of microenvironments affects the migration of single cells, we designed artificial two-state micropatterns consisting of two adhesion sites as previously reported (43). In contrast to the previous study, the micropatterns used here consist of two sites with different geometries, which are connected by a thin bridge. In brief, the surface of the micropatterns is coated with fibronectin, while the surrounding space is passivated using PLL-PEG. As a result, cells only spread within the fibronectin-coated area and remain effectively confined in the micropatterns. We seed MDA-MB-231 human breast carcinoma cells at a concentration low enough to ensure sparse filling of the surface with an average occupancy of one cell per micropattern (Fig. 1B) (for further details, see Material and Methods). After seeding, the cells settle and spread on one of the two square adhesion sites in each pattern. Due to their intrinsic motility, cells then repeatedly transition between the two connected sites in a stochastic manner (Fig. 1A and 1C and Supplementary Movie M1).

**Figure 1.**
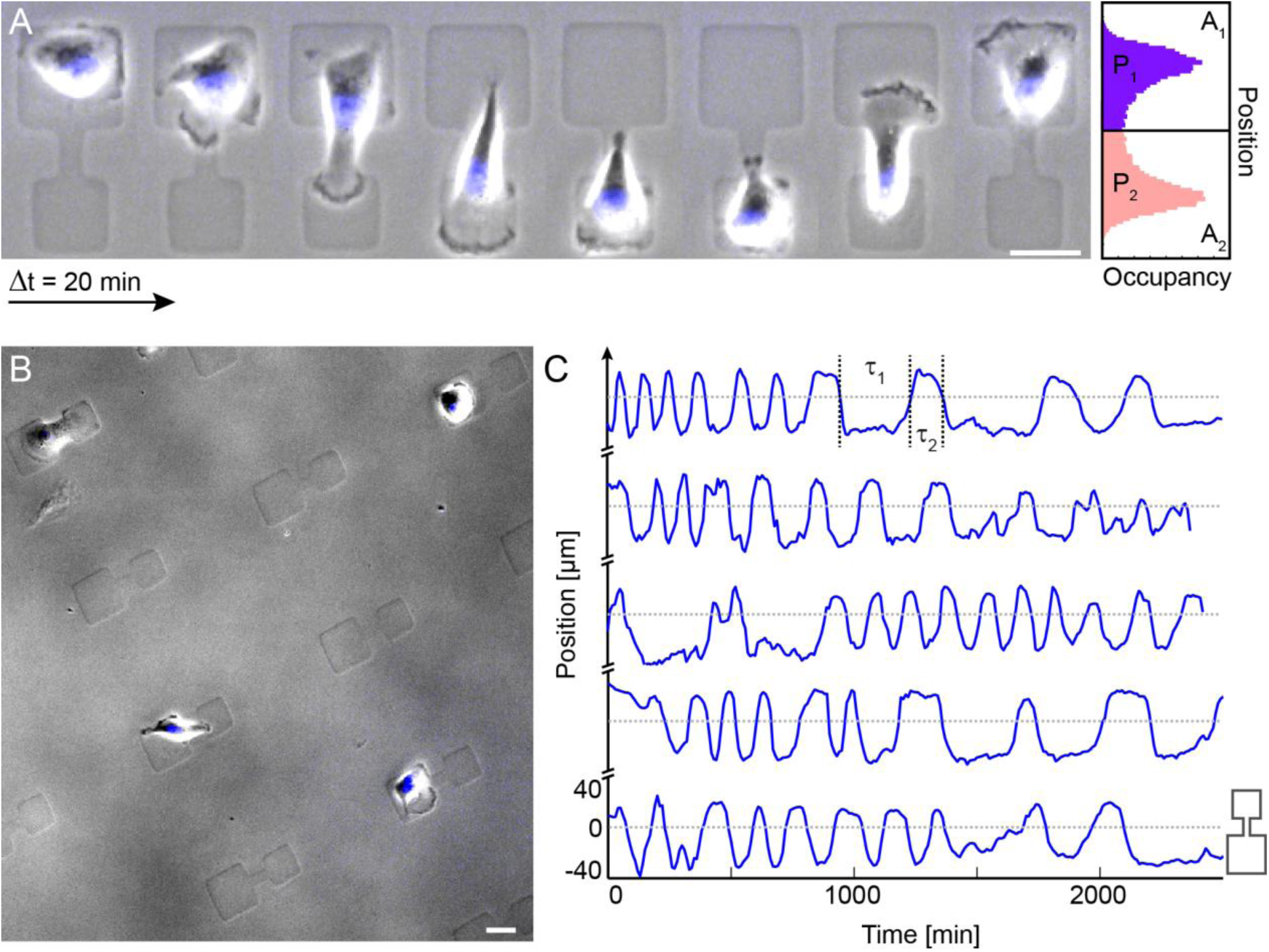
**A**: Time series of a single MDA-MB-231 cell transitioning between two differently sized square adhesion sites. Time intervals between the snapshots are Δt=20 min. The dynamic exploration of the adhesion sites is visible by the extended lamellipodia seen as dark rims along the cell periphery. On the right: Schematic of the probability distribution of the cell nucleus positions along the long axis of the dumbbell. Scale bar: 25 µm. **B**: Overview over experimental setup. Single cells seeded at low density adhere to asymmetric dumbbell-shaped micropatterns. Cell nuclei are stained for semi-automated cell tracking. Scale bar: 25 µm. **C**: Trajectories, as described by the centre-of-mass of the stained cell nucleus, are plotted for several cells on dumbbell patterns of the shown geometry. The definition of dwell times (τ_1_, τ_2_) is shown. Dwell times allow for absolute quantification of cell response.

Transitions between adhesion sites are characterised by a typical order of morphological sequences: Cells that are positioned on one of the two sites exhibit dynamic ruffling of the cell contour and form filopodia, as seen in LifeAct staining (Fig. 4 (ii), Supplementary Movies M8 and M9). Interestingly, short-lived unsuccessful attempts to push lamellipodia beyond the boundaries of the micropattern are clearly visible at the periphery of the cell (black regions in phase contrast images Fig. 1A). In contrast, at the entrance to the bridge, stable lamellipodia form that can proceed to grow across the bridge, which guide the cells towards the other site. Once a lamellipodium reaches the opposite adhesion site, a fan-like broadening of the lamellipodial tip is observed, which is usually followed by the cell body transitioning across the bridge to the other side (Fig. 1A). On the opposite site, the cell spreads into the available adhesion site area and the process repeats.

**Figure 2.**
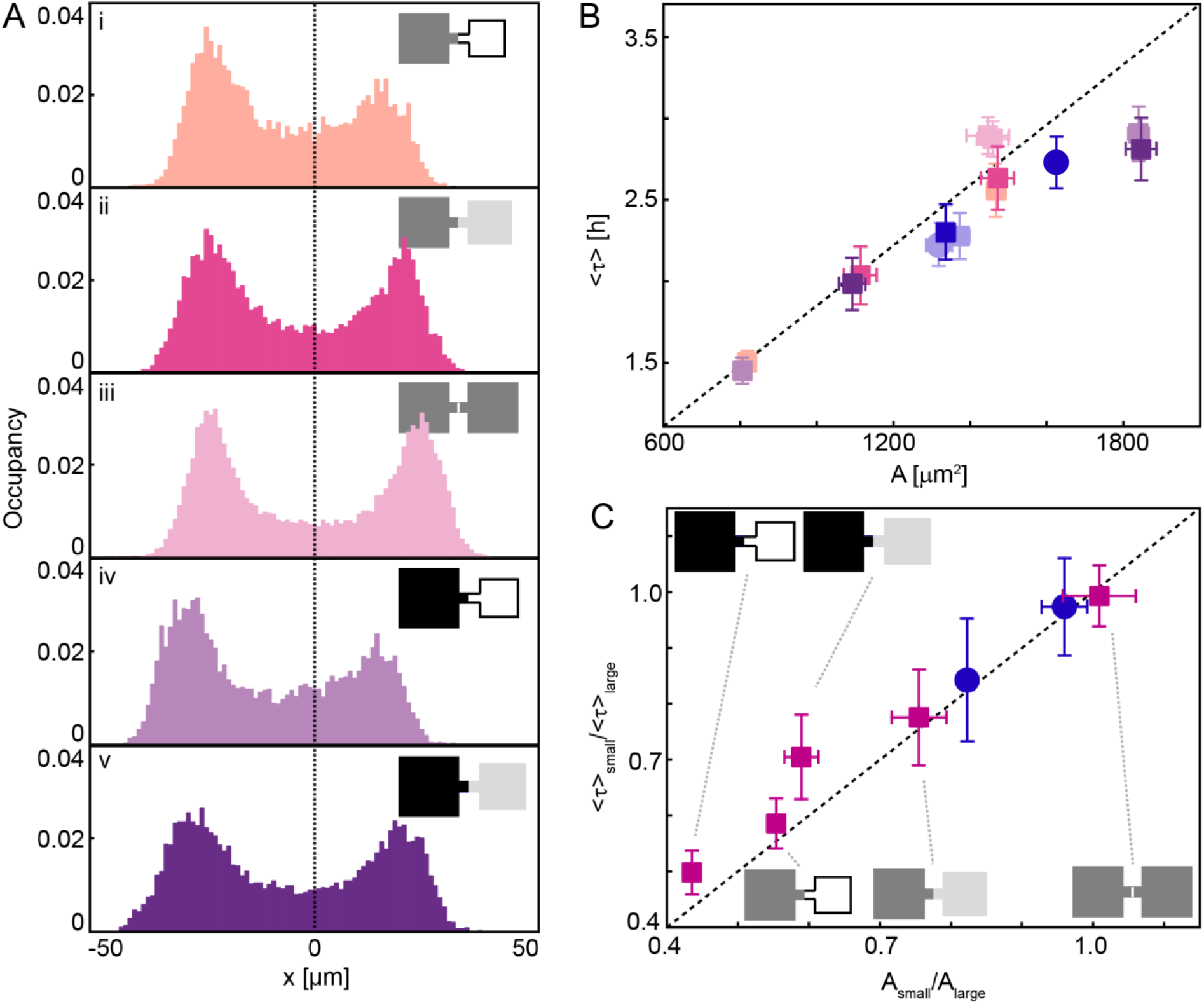
**A**: Occupancies of cells on square adhesion sites with different area. Adhesion sites are depicted in grey shades, same tones denoting the same area. The dotted line represents the middle of the bridge. The occupancies reflect the asymmetry of the underlying micropattern. Third in row (iii) is the symmetric case. (i) and (iii) are adapted from (43). **B**: Mean dwell times, ⟨*τ* ⟩, of cells as a function of square area, A. Dwell times increase with adhesion site area. The dotted line is a guide to the eye. Same colours denote the same two-state geometry, colours correspond to the colours of occupation probability distributions in **A**. Blue circles and squares correspond to dwell times on square-circle micropatterns. Y-Errors are bootstrapping errors, x-errors are standard deviations of weighted area means. **C**: The ratio of mean dwell times plotted against adhesion site area ratio. The dashed line indicates equality. Blue circles correspond to data from square-and circle micropatterns. Y-Errors are bootstrapping errors.

**Figure 3.**
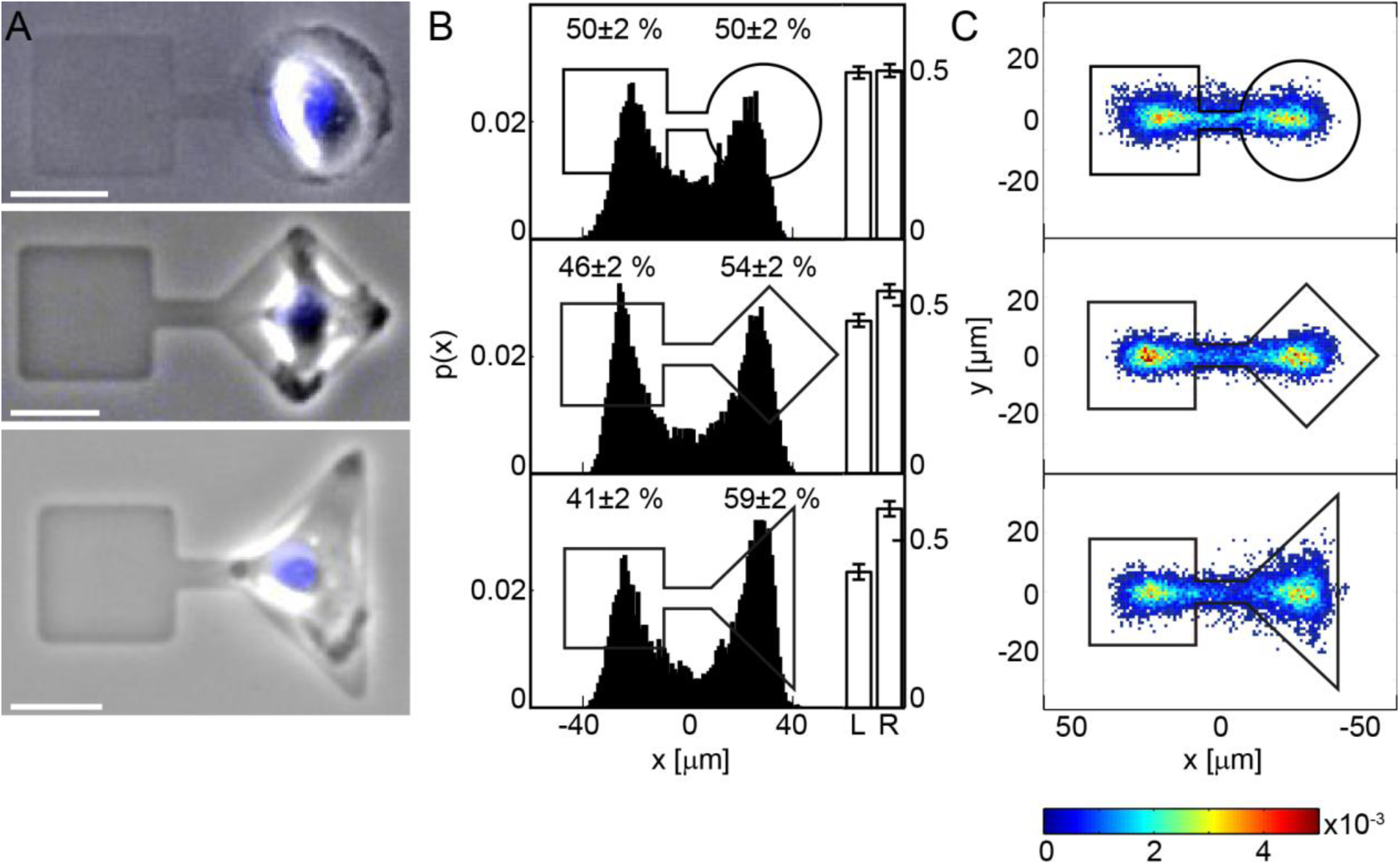
**A**: Snapshots of single MDA-MB-231 cells migrating on two-state patterns with adhesion sites of different geometries and approximately equal area. **B**: Probability distributions of cell positions along the x-axis of the two-state patterns and the corresponding left-right occupation probabilities normalised by the corresponding adhesion site area. An occupation asymmetry is detectable for the square versus rhombus pattern and clearly visible for square versus triangle. Errors are bootstrapping errors. **C**: Two-dimensional probability maps of cell positions with the corresponding micropatterns drawn to scale.

**Figure 4.**
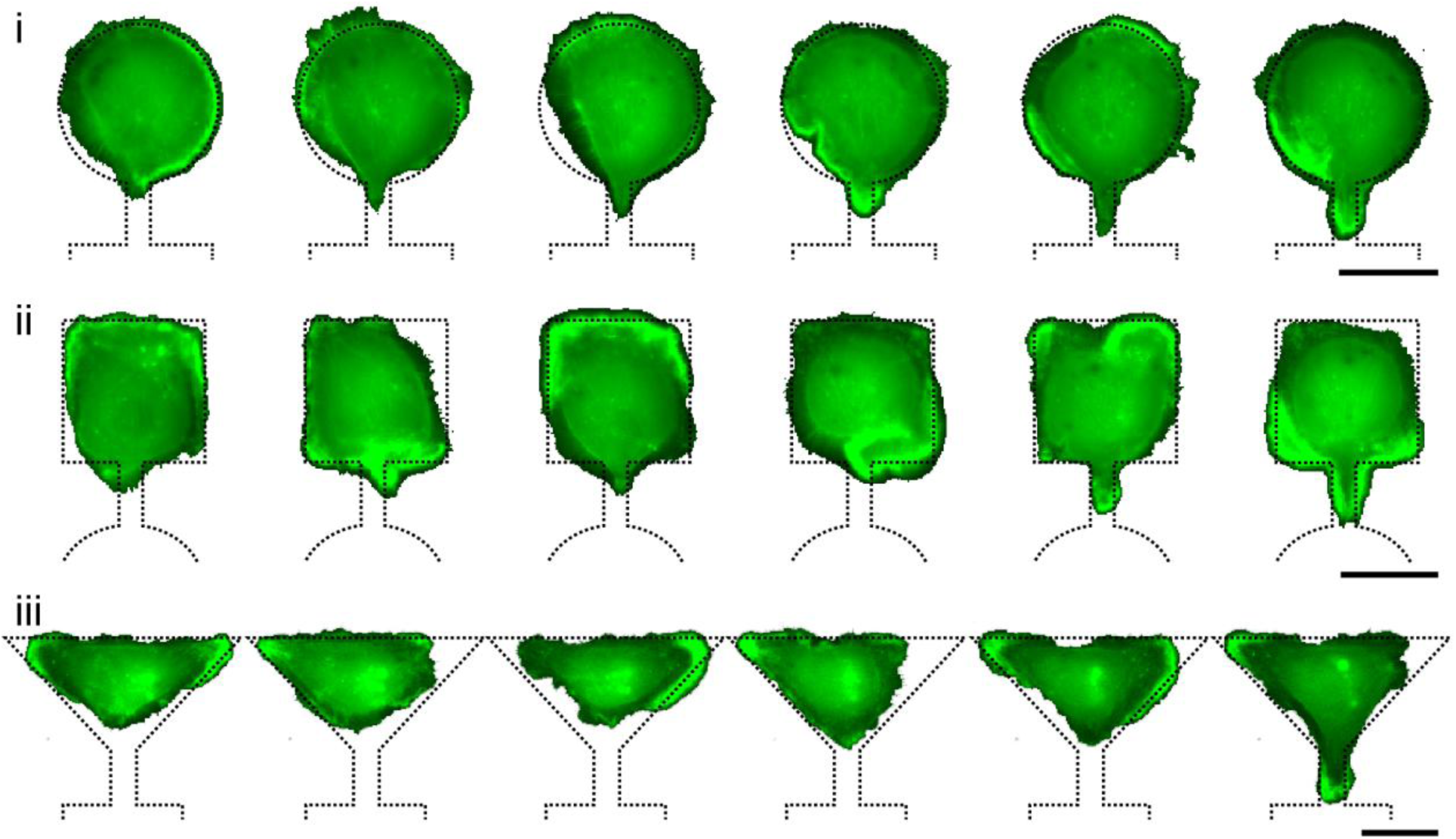
Typical morphological sequences of LifeAct-GFP stained single cells for the asymmetric dumbbells shown in Fig. 3. The distribution of protrusions is seen as bright spots of localized actin along the cell periphery. Actin activity appears more pronounced at adhesion sites’ corners. For improved visibility, cell outlines were selected with the wand tool in Photoshop, and the background colours inverted. Pattern outlines are drawn up to scale. Scale bars: 25 µm.

This cellular hopping process is well captured by the trajectories of the fluorescently-labelled cell nuclei (43). The trajectories span time intervals of up to 50h – their length is limited by cell division or the end of a measurement. Within this time interval, the transition process is stochastic and individual tracks vary from each other. While the cells spend a significant amount of time on the adhesive islands, the transitions themselves occur abruptly (Fig. 1C). These distinct transitions allow an effective reduction of cell positions to two states, separated by the centre of the connecting bridge. A ‘stay’ is defined as the total amount of time the nucleus spends on one side between two crossings of the middle of the bridge. Each of these stays yields a corresponding dwell time, τ.

### Micropatterns with unequally sized adhesion sites exhibit asymmetric occupation probabilities

First, we ask the question how adhesion site area influences the transition statistics. To this end, we perform experiments with square adhesion sites of different edge lengths in the range of 27.2 to 42.3 µm. The connecting bridge has a fixed length of about 16 µm. We determine the occupancy probabilities as a function of position along the main axis of the micropattern with the centre of the bridge defined as *x* = 0. As expected, on equally sized adhesion sites, the occupation probabilities are symmetric (Fig. 2A (iii)). In contrast, the probability distributions corresponding to asymmetric two-state patterns are asymmetric with a clear shift towards higher occupation probabilities on larger adhesion sites (Fig. 2A).

To quantify the dependence of the transition dynamics on the adhesion site area, we plot the mean dwell times ⟨*τ*⟩, averaged over all time traces of the entire ensemble of single cell trajectories, against the corresponding adhesion site areas. We find an almost linear increase of dwell times with indications for a saturation effect for areas larger than approximately 1500 µm^2^. It appears that the dwell times are largely determined by the area of the confining adhesion site. Consistent with this, the dwell times are independent of the size of the opposite adhesion area (Fig. 2B and Fig. S3 in the Supporting Material). In Fig. 2C, the ratio of mean dwell times is plotted against the ratio of adhesion site areas and compared to the direct proportionality relationship between dwell times and adhesion site area (dotted line).

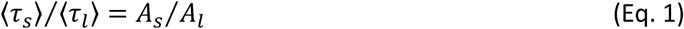

where ⟨*τ*_*s*_⟩, ⟨*τ*_*l*_⟩ denote the mean dwell times and A_s_, A_l_ the area of the small and large adhesion sites, respectively. For area ratios smaller than 0.5, the data slightly deviate from the linear relation. In particular, the two data points with the largest tested adhesion site (A_l_ ≈ 1780 µm^2^) deviate most from linearity (the corresponding dumbbells are shown as inserts with black areas in the left upper part of Fig. 2C). This observation and the saturation effect described in Fig. 2B indicate that the linear relation of dwell times with area is a consequence of confinement. Note that cells only cover the entire adhesion site on average (Supplementary Section 2 and Fig. S1). The mean cell area of unconfined cells is about (940 +/− 37) µm^2^ and hence slightly smaller than the majority of adhesion sites.

It is also noteworthy that for the consideration described above, we can either use the mean values of dwell times or the occupation probabilities. We determined the occupancy probabilities as a function of position along the long axis of the dumbbell (Fig. 2A). The integral of the occupancy probability densities on the left and right sides yield a left versus right occupation ratio, which is equivalent to the ratio of left/right mean dwell times in the limit of long time traces with an equal number of left and right stays. As further detailed in Material and Methods, in a steady-state situation and for sufficiently large statistics, we can equate without further assumptions:

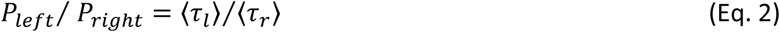

However, while occupation probabilities present a relative measure of preferential cell localisation, the mean dwell times are absolute quantities, which provide a time-scale of the transition dynamics. In the following analysis, we will mostly discuss mean dwell times, which may also be considered as a reverse escape rate, 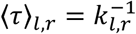, yet without making a statement on the detailed dynamics and nature of the transition mechanism. Taken together, the results so far show that cell occupancies are related to the adhesive area through an approximately linear proportionality relation, and that adhesive area can act as a key control parameter for confined cell dynamics.

### Anisotropic Adhesion Site Shapes Bias Occupancies

We next investigate how the geometrical shape of adhesion sites affects cell migration. To this end, we designed asymmetric two-state micropatterns that connect a square adhesion site to sites with the same area, but a different shape, including circular, rhombical, and triangular sites (Fig. 3). In all cases, the cells perform repeated stochastic transitions, adapt to the shape of the underlying adhesion site and dynamically explore the available area (Fig. 3A and Supplementary Movies M2-M4). To correct for differences in adhesion site areas introduced in the production process, occupation probabilities are normalised by adhesion site areas for square-circle, square-rhombus and square-triangle micropatterns.

Using circles we can test the dependence of dwell times on area or perimeter while keeping the other respective parameter constant. On square-circle micropatterns with equal areas, we observe a symmetric occupation probability distribution (Fig. 3B), and both types of square-circle patterns follow the general trend (blue circles in Fig. 2B, 2C, Fig. S2 and Supplementary Section 3). To relate this finding to the underlying cytoskeletal dynamics driving the migration, we observe LifeAct-GFP transfected cells. Visible as actin hotspots along the cell periphery, a dynamic exploratory motion of the cell within the adhesion sites is apparent. While the distribution of actin appears to be homogenous along the periphery of circular adhesive sites, on square adhesion sites, protrusions form preferentially at the corners. However, overall, very few differences are visible in the actin dynamics, in the type and localisation of actin fibres and locations of protrusion activity between cells adhering to the square and circular adhesion sites (Fig. 4 (i) and (ii) and Supplementary Movie M8).

Rhombical adhesion sites, specifically squares rotated by 45°, allow us to investigate the question whether the corner facing the bridge entrance is able to bias the cell occupancies towards the square site as suggested by previous research (38, 47, 48). Interestingly, the occupation probabilities on square-rhombus patterns show a small bias towards stays on the rhombus (Fig. 3B).

To challenge this unexpected finding, in the next step we designed dumbbells of a square paired with a right-angled triangle. Two-state systems of this kind show a bias towards increased dwell times on the triangular site (Fig. 3B). This finding coincides with the observation that the 2D occupancy distribution on the triangle exhibits a larger spread in the vertical direction, while the spatial distributions on the square, circular and rhombical shapes are very similar (Fig. 3C). In addition, cells on the triangular shape exhibit pronounced protrusion formation at the triangle’s corners (Fig. 3A, Fig. 4 (iii), and Supplementary Movie M9).

Hence, we recapitulate that for sites that are isotropic in the x- and y-directions (square and circle), only adhesive area and not the geometrical shape determines the relative cell occupancy. In contrast, we found that cells take a longer time to escape from rhombical and triangular patterns. Clearly, the triangle has a lower symmetry than the square and circular structures and the protrusions appear to be biased in the vertical direction towards the pointed corners. The rhombus seems to represent an intermediate case, as only a small bias is observed (see also Fig. S3).

### Different orientations of rectangular adhesion sites result in biased occupancies

The observed asymmetry on rhombical and triangular adhesive sites leads to the question of whether adhesion site orientation can determine dwell times. To resolve this question, we designed patterns of equal shape and area but of different orientation. These systems consist of rectangular sites with an aspect ratio of 2:1 in three arrangements: two horizontal rectangles, i.e. with their long axes parallel to the direction of the connecting bridge; two vertical rectangles, i.e. with their long axes perpendicular to the connecting bridge; and a mixed, asymmetric configuration with a horizontal and a vertical rectangle. For the two symmetric arrangements of the rectangular adhesion sites, the observed occupancies are symmetric as expected (Fig. 5B (ii) and (iii)). In contrast, in the case of the asymmetric combination, we find a significant bias towards stays on the upright site (Fig. 5B (i)), supporting our previous assumption that adhesion site anisotropy biases occupancies.

**Figure 5.**
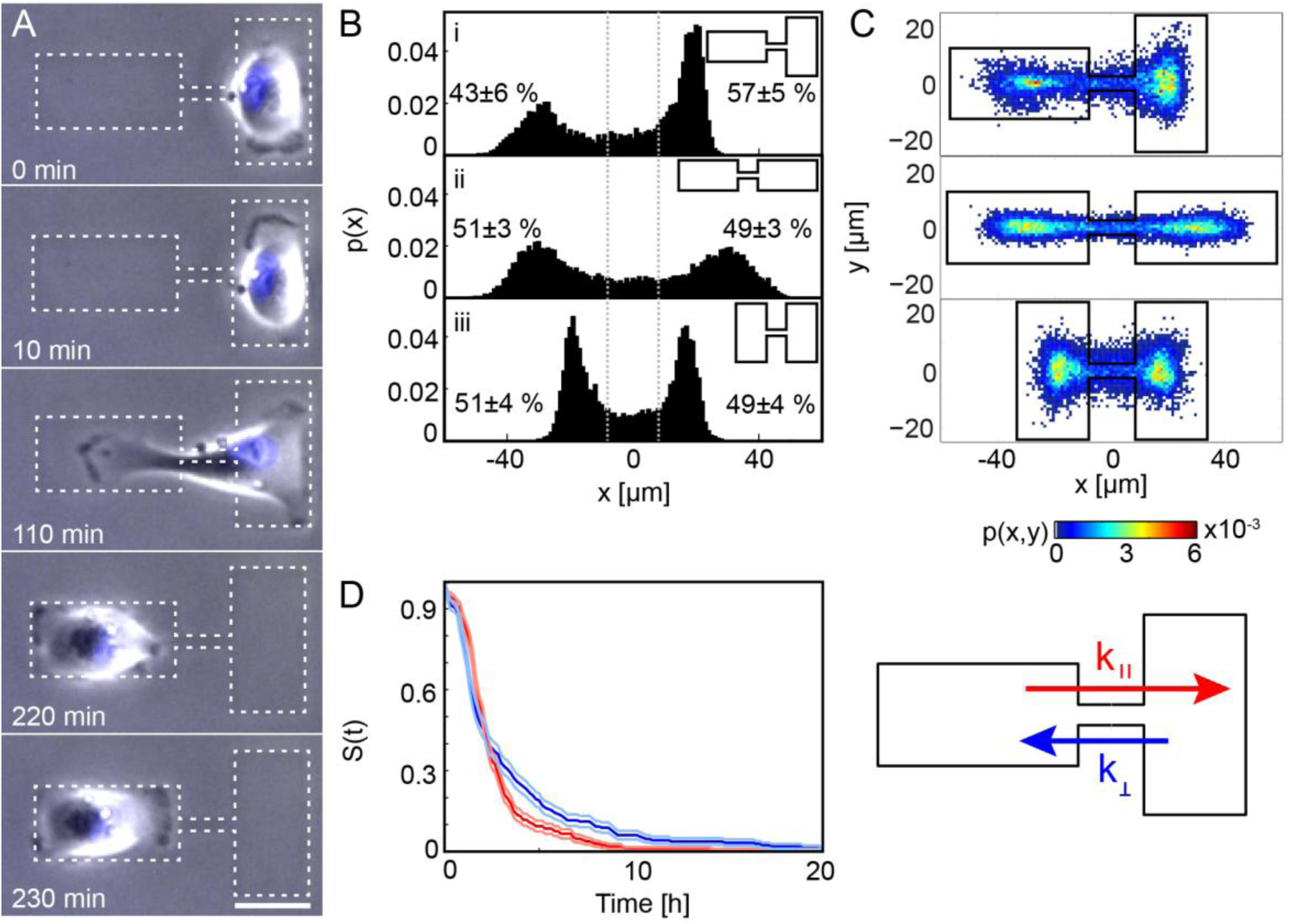
**A**: Time series of a single cell on a rectangular micropattern of orthogonal orientation. The cell preferentially polarizes along the long axes of the rectangles as seen by the formation of lamellipodia in these directions. Scale bar: 25 µm. **B**: The cell probability distribution and left-right occupancies on two-state patterns with equally sized but differently oriented rectangles. The distribution is biased in the case of orthogonal orientation (i). Errors given are bootstrapping errors. **C**: Two-dimensional probability maps of cell positions. A clear difference in the extent of localisation along the y-axis is visible between horizontal and vertical rectangles. **D**: Survival probability functions S(t) of stays in the horizontal and vertical site respectively as indicated in the schematic drawing. Errors are shown as lightly coloured lines. A significant difference in decay behaviour of S(t) is visible for times > 4h.

The observed occupancy asymmetries on the mixed-configuration micropattern can be explained in terms of the steady-state equation Eq. 2, which relates the relative occupancies to the dwell time ratio or, in other words, the ratio of the dynamical escape rates from a horizontal and a vertical state. We observe that on rectangular adhesion sites cells tend to polarize along the long axis of the rectangle (Fig. 5A, 0-10min and 220-230 min, Fig. S5 in the Supporting Material and Supplementary Movies M5-M7), and hence protrusions are primarily observed to form in the direction of the rectangle’s long axis. Thus, one may expect that the escape rate along the long axis is faster than along the short axis, resulting in a biased steady-state distribution.

The full dynamical dependence of the escape process is best represented by the survival probability distribution S(t) which gives the probability that a transition has *not* occurred after time *t* (Fig. 5D). Initially, S(t) for upright rectangles (blue curve) decays faster than S(t) for horizontal rectangles (red curve). However, for stay times longer than 4h, the trend reverses and the blue curve decays significantly slower. This suggests that when transitioning onto the upright rectangle, cells either quickly repolarize or become trapped. The former is possible when cells are reflected by the edge of the micropattern and thus have to turn by 180°. Trapping happens when lamellipodia form along the long axis of the rectangle, making the formation of a lamellipodium along the bridge, a pre-requisite for escape, less likely because the cell is polarized orthogonally to the bridge.

The relation of cell polarization and transition rates is also demonstrated in a more detailed analysis of the two-dimensional cell trajectories. Specifically, we observe a more pronounced motion in the y-direction on vertical rectangles than on horizontal rectangles (Fig. 5C, 6A, Fig. S6). The distribution of velocities in the y-direction is broader on vertical states, indicating that the cells polarize and accelerate along the long axis of the rectangle, orthogonal to the axis of the micropattern (see also Supplementary Section 5, Fig. S7). Moreover, we find that whenever there is pronounced displacement in the y-direction, the extent of motion in the x-direction decreases (Fig. 6B). This suggests that the cell polarization in one direction suppresses motion in the orthogonal direction. Thus, escape rates from vertical rectangles are reduced by the induced polarization orthogonal to the direction of the transitions. For a more detailed discussion of escape times on rectangular dumbbell micropatterns we refer to the Supplementary Section 5.

**Figure 6.**
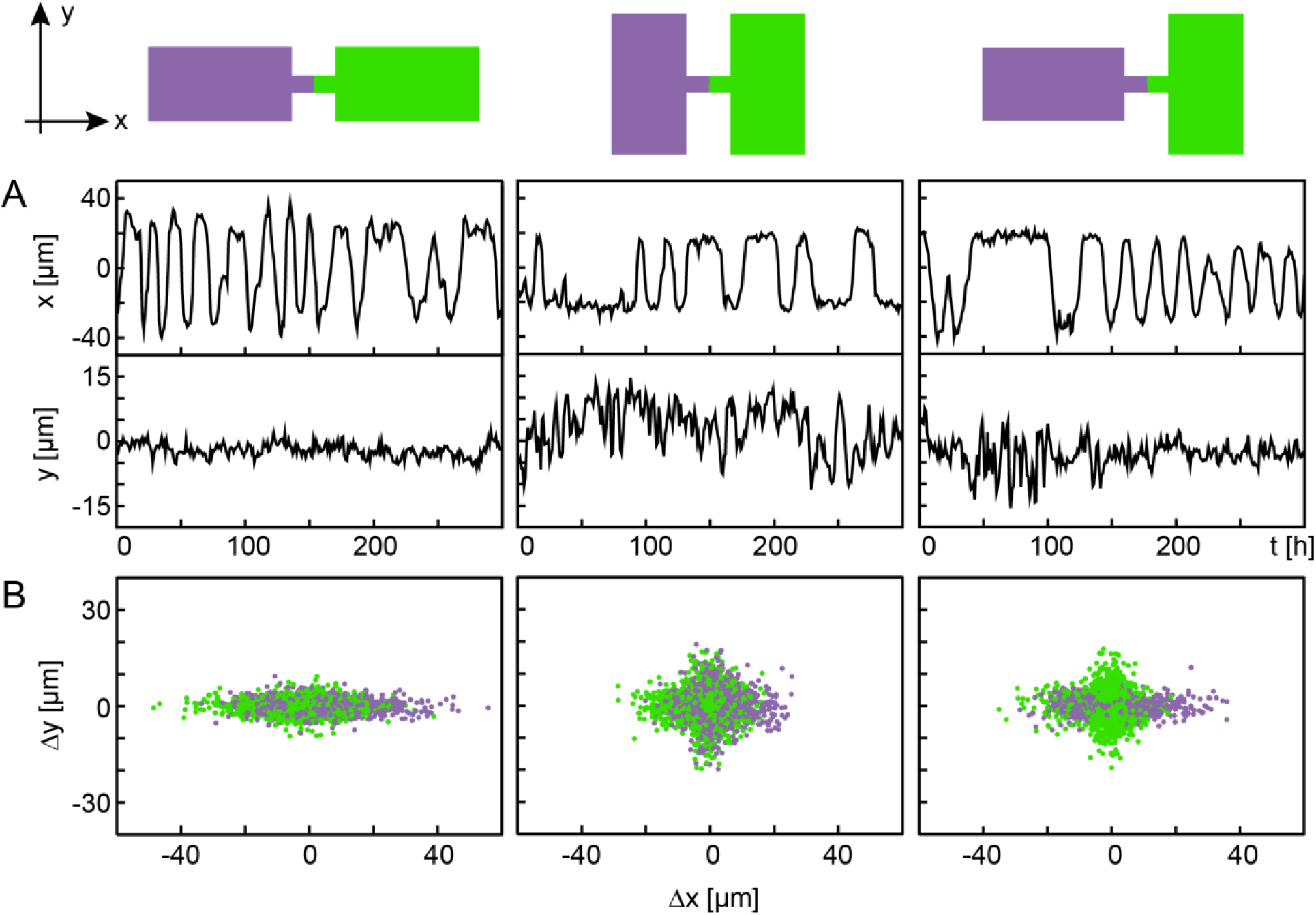
**A**: Comparison between x- and y-motion: Exemplary single-cell trajectories in x- and y-direction for each of the three rectangle setups. The displacement fluctuations along the y-axis are larger for vertical rectangles than for horizontal rectangles. **B**: Displacements in x- and y-direction with respect to the centres of the rectangles. The colours of the dots correspond to the left or right adhesion site, as indicated in the schematic drawings. It is clearly visible that where the extent of x-motion is large, motion in the y-direction is limited, and vice versa.

In summary, we found that the orientation of anisotropic adhesion sites can bias cell occupancies since migrating cells tend to polarise along the long axis of these sites, leading to a reduced escape rate when this axis is orthogonal to the transition axis.

## Discussion

In this work, we studied cell migration on artificial two-state micropatterns, which allow for a statistical analysis of the relative preference of cells for two opposing adhesion sites with different geometric properties. Occupancies are described in terms of relative occupation probabilities or absolute mean dwell times. We observed a preferred occupancy with increasing area in the case of two square adhesion sites. The finding is in agreement with previously reported ‘dimension sensing’ of cells on thin 1D lines interspersed with wider rectangles (36). Larger adhesive areas induce longer dwell times, which is in qualitative agreement with the observation that freely moving cells exhibit reduced cell motility with larger spreading area (49, 50). Also, recently, Guo et al. reported that a large cell adhesion area is linked to smaller cell volume and an increase in cell stiffness due to water efflux in a spread out state (51). A stiffer cell cortex is likely associated with slower migration (52). Thus, a potential explanation for the area dependence of escape dynamics (Fig. 2B and 2C) might be a transient stiffening of the cell cortex on sites with larger adhesion area.

We found only a weak dependence of occupation probabilities on shape in sites that are symmetric under 90 degree rotations. In particular, cells on circular, square and rhombical adhesion sites exhibit a very similar transition behaviour (Fig. 2B, 2C, Supplementary Section 4, Fig. S3). This is remarkable since some of these sites have different perimeters, which is a parameter that frequently enters elastic cell models which assume a global line tension and hence a dependence on perimeter (53, 54). While previous studies suggest a difference in the localized distribution of cellular protrusions on square and circular micropatterns (28, 55), we did not observe any influence of square or circular shape on occupancies, which should be sensitive to the likelihood of directional protrusions. This finding, however, is consistent with the reported lack of cell polarity on square and circle micropatterns (28). The absence of induced cell polarisation is also demonstrated in experiments, where cells showed no preferred direction of motion when released from squares or circular micropatterns by removal of a micropatterned frame (29).

The role of acute corners and stress fibres on the transition dynamics needs further inspection. It was reported that stress fibres running along straight edges impede protrusion formation, while stress fibres meeting in corners are easier to penetrate (38, 47, 48). Intriguingly, in the presence of other guidance cues such as corners located orthogonally to the axis of transitions, we did not observe increased transition rates where corners were pointed towards the connecting bridge. Specifically, the presence of acute angles in triangles seems to stretch and orient cells in a direction perpendicular to the transition axis. This could be related to the increased likelihood of cells to extend lamellipodia from acute angles (30). Also, the importance of pattern orientation became clearly visible in the rectangular system where a directed formation of lamellipodia orthogonal to the direction of transitions between the adhesion sites is achieved, and therefore the motion becomes anisotropic (28, 29). It needs to be emphasized, however, that our measurements on geometrical effects are dynamic and deviations from observations of static cell patterns may be a matter of the time-scales of actin cytoskeleton remodelling. While the steady-state probability distribution is insensitive to variations of isotropic shapes, we cannot exclude that dynamic features will be affected. For the case of symmetric two-state micropatterns, we found non-linear stochastic dynamics with characteristic limit cycle or bistable behaviour (43). In future theoretical work, we will deepen the insights into the geometry-dependence of the migration dynamics and the corresponding deterministic and stochastic dynamics.

## Conclusion

In summary, we created asymmetric two-state micropatterns, which allow for a quantitative assessment of geometric determinants in confined cell migration. The comparison of mean dwell times and steady-state occupancy probabilities in different adhesive geometries identifies relevant factors, such as adhesion site area and geometry, and quantifies their effect on cell migration. The statistics of individual trajectories and transition rates also suggest that escape rates could be an indirect measure of protrusion dynamics in confining microenvironments. In general, single-cell microarrays enable a highly parallel assessment of an ensemble of cells in defined boundaries. Two-state microarrays in particular could be extended to probe cellular affinities for protein-coated surfaces by creating patterns with different chemical functionalities. In addition, as the two-state micropatterns allow for repeated observation of transition events of individual cells, the leading-edge dynamics could be studied under defined boundary conditions and with high statistical certitude. The forced transitions in artificial microenvironments could also, in future work, be related to relevant, disease-related states of cells like deformability (56) and, for example, serve to characterize migratory phenotypes such as invasiveness (57). Hence, two-state micropatterns represent a unique single-cell migration assay with the capacity to assess relative preferences of cells for microenvironments and to define novel metrics for cell-migration phenotypes.

## Author contributions

A.F., C.S., P.J.F.R. and J.O.R designed experiments; A.F. performed experiments; A.F. and D.B.B. analyzed data. A.F., D.B.B., C.S., C.P.B. and J.O.R. interpreted the experiments and wrote the manuscript.

## Acknowledgments

Financial support of the German Science Foundation (DFG) for the collaborative research center SFB 1032 project B01 and B12 is acknowledged. D.B.B. is supported by a DFG fellowship within the Graduate School of Quantitative Biosciences Munich and by the Joachim Herz Stiftung. We thank C. Leu for the preparation of wafers, A. Reiser for providing the transfection protocol, G. Schwake for preparation of the LifeAct-mRNA, S. Reinhardt for measuring cell areas and E. Petrov for fruitful discussions.

