## Supplementary Information for "Area and Geometry Dependence of Cell Migration in Asymmetric Two-State Micropatterns"

**Contents:**

0. Movie Description

1. Additional Materials and Methods

1.1. Detailed Micropattern Dimensions

1.2. Cell Exclusion Criteria

1.3. Cell Area Determination

2. Cell Areas on Square Adhesion Sites

3. Square-Circle Pattern of Equal Perimeter

4. Survival Functions on Adhesion Sites of Different Shapes

5. Rectangle Micropatterns

5.1. Survival Functions as a Function of Bridge Length

5.2. Migration Behaviour on Rectangle Micropatterns

#### 0. Movie Description

**Supplementary Movie M1:**

Single MDA-MB-231 cell migrating repeatedly back and forth between two square adhesion sites of different areas ($A_{left}$=1405 µ$m^2$ and $A_{right}$=755 µ$m^2$). The nucleus is stained with Hoechst 33342 to allow semi-automated cell tracking.

**Supplementary Movies M2-M4:**

Single MDA-MB-231 cells migrating repeatedly back and forth between two adhesion sites of comparable areas but of different shapes: square and circle, square and rhombus and square and triangle.

**Supplementary Movies M5-M7:**

Single MDA-MB-231 cells migrating on rectangle micropatterns of different orientations. It is visible how the cells polarize along the long axes of the rectangles. Bridge length=16.2 µm.

**Supplementary Movies M8-M9**

LifeAct-GFP stained single MDA-MB-231 cells migrating on square-circle (equal areas) or square-triangle micropatterns. Actin hotspots along the cell periphery are visible which are preferentially localized in the patterns' corners.

### 1. Additional Materials and Methods

#### 1.1. Detailed Micropattern Dimensions

| x-dimensions left site [µm]* | x-dimensions right site [µm]* | Area left [$\mu m^2$] | Area right [$\mu m^2$] | Further information | Number of cells analysed | Bridge length x width [µm] |
|---|---|---|---|---|---|---|
| 37.5 ± 0.3 | 27.5 ± 0.2 | 1405 ± 22.5 | 754.7 ± 11.0 | Square-square | 47 | (16.0 ± 0.3) x (7.8 ± 0.3) |
| 37.6 ± 0.6 | 32.4 ± 0.7 | 1410.6 ± 45.1 | 1052 ± 45.4 | Square-square | 68 | |
| 37.2 ± 0.8 | 37.4 ± 0.6 | 1385.1 ± 59.5 | 1397.3 ± 44.9 | Square-square (symmetric) | 169 | |
| 42. 1 ± 0.3 | 27.3 ± 0.4 | 1776.2 ± 25.3 | 743.1 ± 21.8 | Square-square | 58 | |
| 42.2 ± 0.5 | 32.1 ± 0.6 | 1782.1 ± 42.2 | 1028.8 ± 38.5 | Square-square | 74 | |
| 36.5 ± 0.4 | 40.3 ± 0.6 | 1329.7 ± 29.2 | 1275 ± 38.0 | Square-circle (equal area) | 67 | (14.5 ± 0.2) x (6.1 ± 0.1) |
| 35.9 ± 0.2 | 44.6 ± 0.3 | 1291.9 ± 14.4 | 1580.2 ± 21.0 | Square-circle (equal perimeter) | 98 | |
| 37.6 ± 0.2 | 46.7 ± 0.5<br>49.6 ± 0.6 | 1413.8 ± 14.3 | 1243.2 ± 24.7 | Square-rhombus | 63 | (17.7 ± 0.1) x (7.7 ± 0.1) |
| 37.8 ± 0.2 | 33.5 ± 0.3<br>67.7 ± 0.7 | 1428.8 ± 17.3 | 1263.8 ± 16.5 | Square-triangle | 82 | |
| 48.9 ± 0.7<br>24.9 ± 0.4 | 48.9 ± 0.8<br>24.9 ± 0.3 | 1217.6 ± 26.2 | 1217.6 ± 24.6 | Lying rectangles | 78 | (16.2 ± 0.1) x (5.0 ± 0.2) |
| 24.9 ± 0.4<br>47.9 ± 0.5 | 24.8 ± 0.3<br>47.8 ± 0.4 | 1192.7 ± 22.1 | 1185.4 ± 15.3 | Upright rectangles | 65 | |
| 48.2 ± 0.7<br>24.3 ± 0.4 | 25.2 ± 0.2<br>47.8 ± 0.5 | 1171.3 ± 24.0 | 1204.6 ± 16.9 | Mixed rectangles | 59 | |

**Table S1.** Measures of all used dumbbell geometries, calculated adhesion site areas and numbers of cells analysed for each geometry. Orange shading denotes adhesion sites designed to have comparable areas. Errors are weighted standard deviations as different statistics were gathered from each experiment and an experiment-to-experiment variation between pattern dimensions is unavoidable. * Y-dimensions of longest axis are given for anisotropic shapes (bottom numbers).

The following channel lengths were probed with different rectangle orientations: (8.2 ± 0.3) µm, (16.2 ± 0.1) µm (as presented in the main text), (24.8 ± 0.2) µm, and (34.7 ± 0.1) µm.

#### 1.2. Cell Exclusion Criteria

To limit the effects of abnormal migration behaviour, we deem cells suitable for further analysis if they comply with the following criteria (1):

1. Only a single cell occupies the micropattern. Observations are stopped when the cell rounds up for division.
2. We only include trajectories of cells which, including their protrusions, are entirely confined within the borders of the micropattern.
3. Transition statistics are only included after the first and last observed transition. This avoids start- and end-of-measurement artefacts in determining dwell times.
4. The cell shows no abnormalities such as multiple nuclei or the occurrence of cell death at any time during the whole experiment.
5. In the vast majority of cases the cell performs complete transitions. A complete transition is defined by the fact that no parts of the cell adhere to the previous adhesion site once the nucleus has entered the new adhesion site.

Criteria 1-4 are basic requirements for single cell studies on micropatterns. Criterion 5 is specific to our dumbbell setup and, depending on the cell type, may be a rather strong constraint. We previously found that criterion 5 can be relaxed without affecting general results (1).

##### 1.3. Cell Area Determination

Cell areas were determined manually with ImageJ's Ivussnakes Plugin (2) by tracing the cell outlines visible in the phase contrast images. Cell areas were determined for cells complying with the above exclusion criteria and only at times where the cell was fully confined (i.e. where there were no protrusions inside the channel). As this approach overestimates the influence of smaller cells, which are naturally better confined, a mean cell area was calculated for every stay (in contrast to calculating a mean over all frames used) and an average of these areas was taken as final cell area.

#### 2. Cell Areas on Square Adhesion Sites

To test the influence of cell area on the mean dwell times, we measure the average cell areas that cells actually cover on the differently sized square adhesion sites. In Fig. S1, the dependence of cell area on available adhesion site area is shown. It is clearly visible that only for the two smallest adhesion sites cells fully occupy the adhesion site, and thereby reflect in their area the area of the adhesion sites. For larger adhesion sites, the cell area still increases but does so more slowly. This means that on large adhesion sites, cells frequently do not fully fill all available area of the confining site.

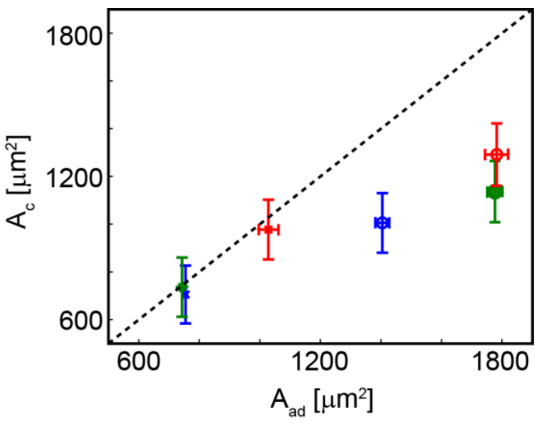


**Figure S1**. Cell areas $A_c$ on adhesion sites of different areas $A_{ad}$. The cell area increases with provided adhesion area. The increase in cell area is not equal to the increase of provided adhesion areas. The dashed line is a guide to the eye corresponding to $A_c = A_{ad}$. Same colours denote cell areas taken from the same dumbbell geometry. Error bars of the cell area are obtained by bootstrapping.

#### 3. Square-Circle Pattern of Equal Perimeter

To probe the influence of shape on cellular dwell times, we have created square-circle micropatterns with adhesion sites of either equal area (data shown in main text) or the same perimeter (Fig. S2). We find that occupation probabilities on square-circle patterns are biased towards the circle if the square and circular adhesion sites have the same perimeters. When the corresponding mean dwell times are added to the plot of mean dwell times against adhesion site areas, the data points follow the same linear dependence on area as seen for square-only micropatterns (Fig. 2B and 2C).

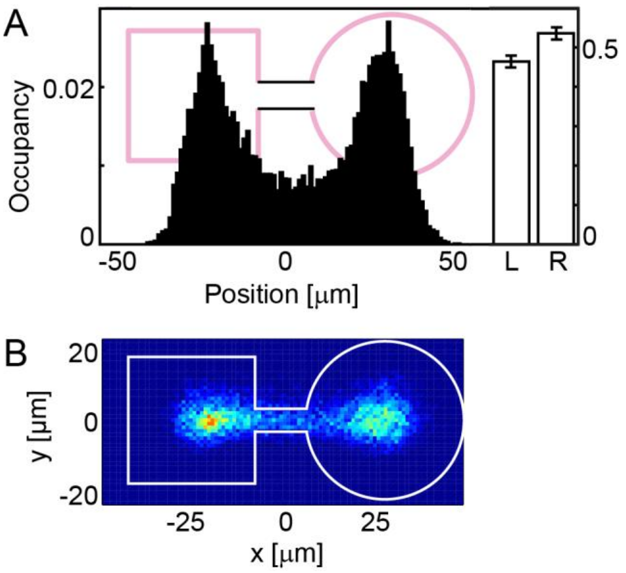


**Figure S2. A**: Probability distribution in x-direction of cell positions on square-circle micropattern with adhesion sites of equal perimeter. A significant asymmetry of stay probabilities is visible, with stays biased towards the circular adhesion site. **B**: 2D probability distribution of cell positions on the same geometry. A broader spread of positions is visible on the circular site.

#### 4. Survival Functions on Adhesion Sites of Different Shapes

We measured the dwell times in repeated experiments for square, circular, rhombical and triangular adhesion sites of similar areas. While for the circle the survival function of stay times, S(t), does not differ from those measured on square adhesion sites, a gradual broadening of S(t) is visible for the rhombical and triangular geometry (Fig. S3).

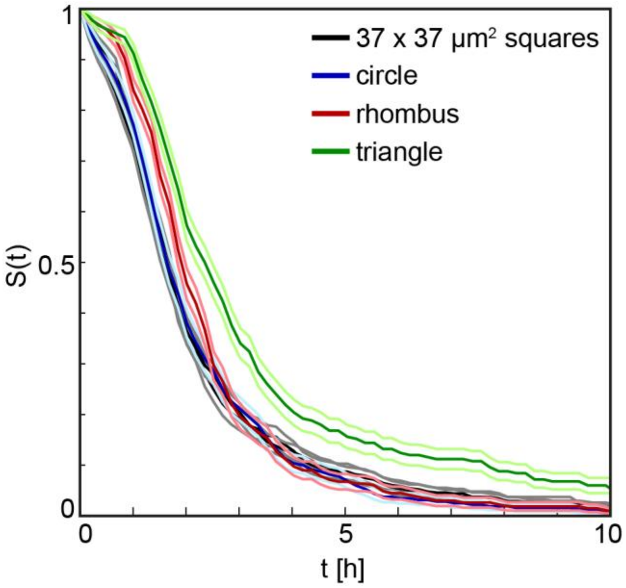


**Figure S3**. Variability of survival functions S(t) for adhesion sites of approximately the same area. S(t) for stays on square adhesion sites are plotted in black. In blue, the survival function for stays on circular adhesion sites with the same area is shown. The red curve shows S(t) for the rotated square (rhombus), the green curve shows S(t) for triangular adhesion sites. Bootstrapping errors are plotted in light colours.

#### 5. Rectangle Micropatterns

##### 5.1. Survival Functions as a Function of Bridge Length

To further explore the migration of cells between anisotropic states, we also vary the length of the connecting bridge between the two rectangles. Due to their identical orientation, we expect the survival probability functions on the symmetric upright rectangle dumbbell and on the upright rectangle from the mixed orientation setup to be identical. The same should be true for horizontal rectangles. From Fig. S4A, where survival probability functions for corresponding rectangle orientations are shown in the same plot and for different channel lengths, it is evident that cells behave differently for the same rectangle orientations, depending on the overall composition of the two-state system. Furthermore, we observe an increase of the mean dwell times with channel length for all rectangle orientations and setups. This behaviour is in qualitative agreement with our observations on symmetric square systems (1). Strikingly, the dwell times on upright rectangles in the mixed setup are generally longer than the corresponding stay times on upright rectangles in the symmetric setup. Furthermore, there is no significant difference in stay times between the horizontal rectangles and the upright rectangles from the mixed orientation setup (Fig. S4B).

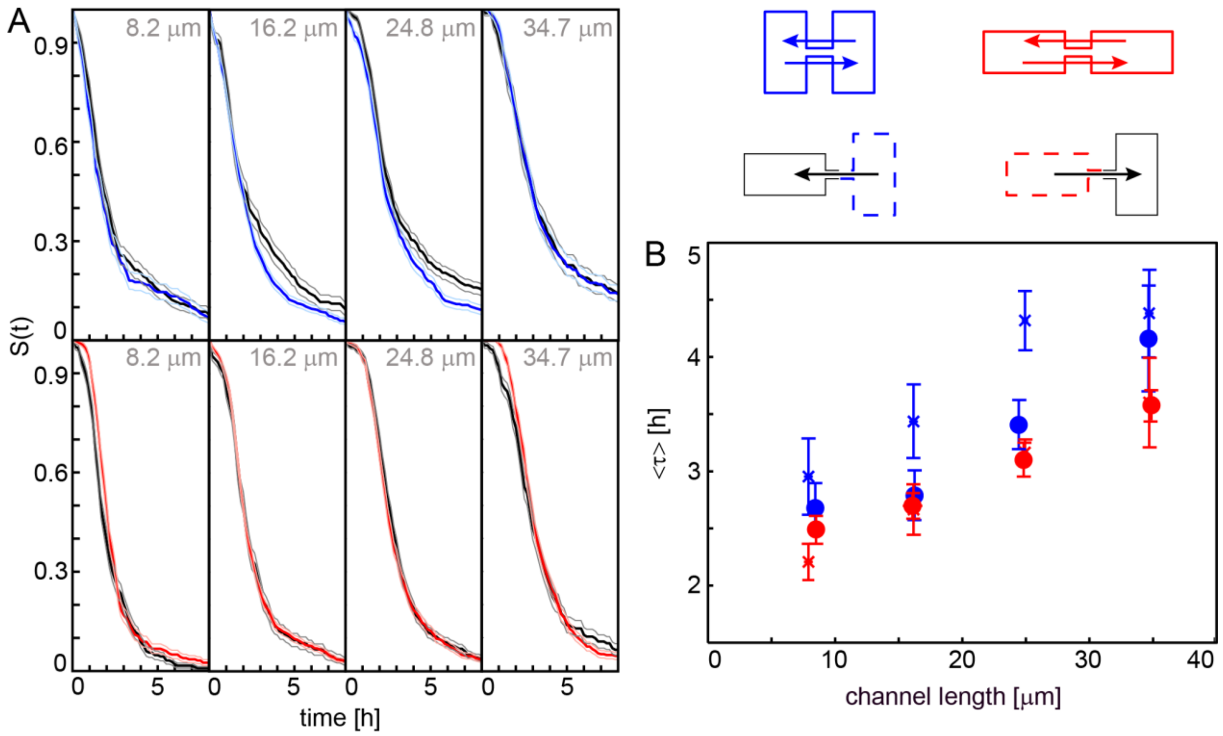


**Figure S4**. **A**: Survival functions S(t) for different channel lengths. In each plot, the survival function is shown for rectangles of the same orientation but from different setups, the black line always corresponding to mixed-orientation dumbbells. Despite the rectangles having the same orientation, deviations in the decay behaviour of S(t) are visible for longer times (t>3h) for upright rectangles connected by channels of medium lengths. In contrast, some differences in S(t) are visible for horizontally-oriented rectangles (lower plots) for short stay times. **B**: Mean stay time <τ > plotted against channel length for all tested rectangle orientations. An approximately linear increase of mean dwell times with increasing channel length is visible for all orientations. The blue cross datapoints correspond to stay times measured on the upright rectangle, in the mixed orientation setup, the solid blue circle data correspond to the symmetric setup of upright rectangles, the red crosses corresponds to data from the lying rectangle of a mixed-orientation setup and the solid red circles correspond to stay times measured on dumbbells with equally oriented, horizontal rectangular adhesion sites.

##### 5.2. Migration Behaviour on Rectangle Micropatterns

In all rectangle orientations, cells are observed to form protrusions along the long axis of the rectangle (Fig. S5). This results in a broadening of the probability distribution of y-positions on the vertical rectangles relative to the horizontal case (Fig. S6). This is the case in all vertical rectangles, both in the symmetric and the mixed setup. The anisotropies also significantly affect the probability distributions of velocities, which are always broader along the long axis of each rectangle (Fig. S7). Taken together, these results show that there is a significant difference in migratory activity along the long and short axes of the rectangles.

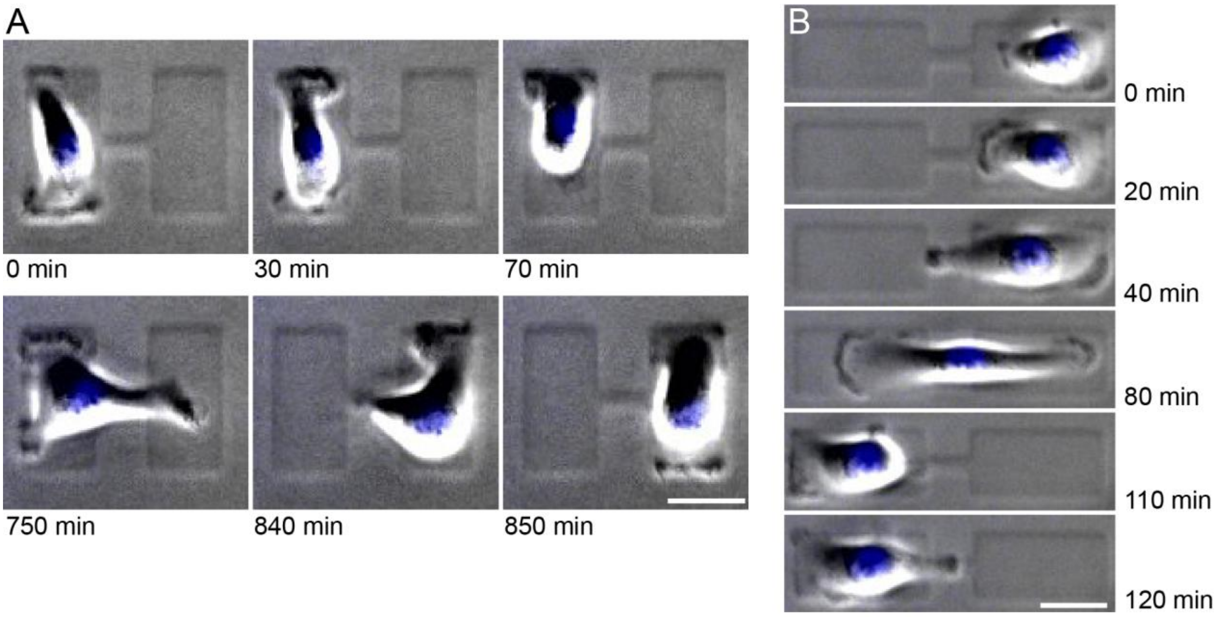


**Figure S5. A**: Time series of a single cell migrating on a dumbbell made of vertical rectangles (bridge length L = 16.2 µm). It can be seen how lamellipodia are formed along the long axis of the rectangle. Scale bar: 25 µm. **B**: Time series of a single cell transitioning on horizontal rectangles. The cell is seen to migrate within each rectangle until it meets the dumbbell border. Scale bar: 25 µm.

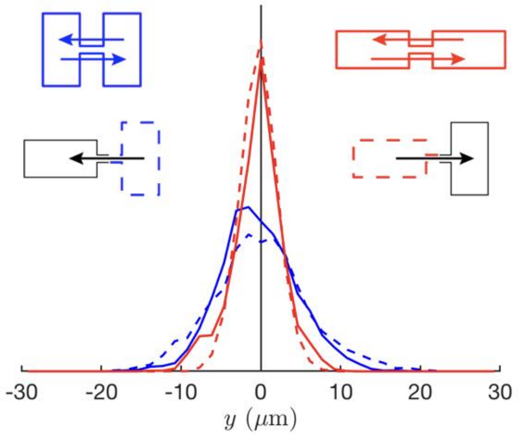


**Figure S6.** Probability distributions of positions in the vertical direction on rectangle micropatterns with bridge length L=16.2 µm. The cartoons serve as a legend to the four distributions.

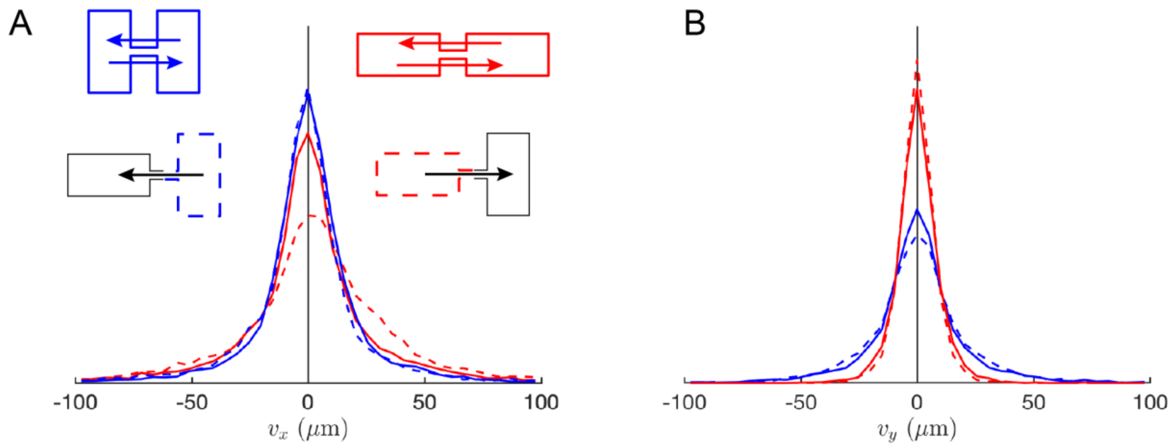


**Figure S7.** Probability distributions of velocities in horizontal and vertical directions (bridge length L = 16.2 µm). **A**: Distribution of velocities along the micropattern's main direction (x-direction). **B**: Distribution of velocities in the direction orthogonal to the main axis of the micropattern (y-direction). The colour code is indicated by the cartoons.
